# An ancient truncated duplication of the anti-Mullerian hormone receptor type 2 gene is a potential conserved master sex determinant in the Pangasiidae catfish family

**DOI:** 10.1101/2022.01.14.475871

**Authors:** Ming Wen, Qiaowei Pan, Elodie Jouanno, Jerome Montfort, Margot Zahm, Cédric Cabau, Christophe Klopp, Carole Iampietro, Céline Roques, Olivier Bouchez, Adrien Castinel, Cécile Donnadieu, Hugues Parrinello, Charles Poncet, Elodie Belmonte, Véronique Gautier, Jean-Christophe Avarre, Remi Dugue, Rudhy Gustiano, Trần Thị Thúy Hà, Marc Campet, Kednapat Sriphairoj, Josiane Ribolli, Fernanda L. de Almeida, Thomas Desvignes, John H. Postlethwait, Christabel Floi Bucao, Marc Robinson-Rechavi, Julien Bobe, Amaury Herpin, Yann Guiguen

## Abstract

The evolution of sex determination (SD) mechanisms in teleost fishes is amazingly dynamic, as reflected by the variety of different master sex-determining genes identified, even sometimes among closely related species. Pangasiids are a group of economically important catfishes in many South-Asian countries, but little is known about their sex determination system. Here, we generated novel genomic resources for 12 Pangasiid species and provided a first characterization of their SD system. Based on an Oxford Nanopore long-read chromosome-scale high quality genome assembly of the striped catfish *Pangasianodon hypophthalmus*, we identified a duplication of the anti-Müllerian hormone receptor type II gene (*amhr2*), which was further characterized as being sex-linked in males and expressed only in testicular samples. These first results point to a male-specific duplication on the Y chromosome (*amhr2by*) of the autosomal *amhr2a*. Sequence annotation revealed that the *P. hypophthalmus* Amhr2by is truncated in its N-terminal domain, lacking the cysteine-rich extracellular part of the receptor that is crucial for ligand binding, suggesting a potential route for its neofunctionalization. Short-read genome sequencing and reference-guided assembly of 11 additional Pangasiid species, along with sex-linkage studies, revealed that this truncated *amhr2by* duplication is also conserved as a male-specific gene in many Pangasiids. Reconstructions of the *amhr2* phylogeny suggested that *amhr2by* arose from an ancient duplication / insertion event at the root of the Siluroidei radiation that is dated around 100 million years ago. Altogether these results bring multiple lines of evidence supporting that *amhr2by* is an ancient and conserved master sex-determining gene in Pangasiid catfishes, a finding that highlights the recurrent usage of the transforming growth factor β pathway in teleost sex determination and brings another empirical case towards the understanding of the dynamics or stability of sex determination systems.

## INTRODUCTION

Catfishes (Order Siluriformes) with approximately 4,000 species (Sullivan, Lundberg, & Hardman, 2006) are economically and ecologically important fish worldwide. Among catfishes, the Pangasiid family (Pangasiidae) is recognized as a monophyletic group including four extant genera, i.e., *Helicophagus, Pangasianodon, Pangasius* and *Pteropangasius* (Pouyaud, Gustiano, & Teugels, 2016). These species have a wide range of habitats both in fresh and brackish water across southern Asia, from Pakistan to Borneo (Roberts & Vidthayanon, 1991). Many Pangasiids, because of their rapid growth rate, are also important aquaculture species, such as *Pangasius bocourti*, *Pangasius djambal* and *Pangasianodon hypophthalmus* (Lazard, Cacot, Slembrouck, & Legendre, 2009). *P. hypophthalmus*, for example, has become a major aquaculture species extensively farmed in many Asian countries (Anka, Faruk, Hasan, & Azad, 2014; Na-Nakorn & Moeikum, 2009, p.; Phuong & Oanh, 2010; Singh & Lakra, 2012) and has even been recently introduced in the Brazilian finfish aquaculture.

Sex determination (SD) mechanisms have not been investigated in detail in Pangasiid catfishes, but genetic sex-linked markers that could facilitate broodstock management for aquaculture or conservation purposes, have been searched without success in both *P. hypophthalmus* and *P. gigas* (Sriphairoj, Na-Nakorn, Brunelli, & Thorgaard, 2007). SD in vertebrates can rely on genetic (GSD for genetic SD), environmental (ESD for environmental SD) or a combination of both genetic and environmental factors (such as thermal effects on GSD = GSD+TE) (Baroiller, D’Cotta, & Saillant, 2009; Kobayashi, Nagahama, & Nakamura, 2013; Ospina-Alvarez & Piferrer, 2008). In teleost fishes, SD has been found to be extremely plastic with both GSD, ESD and GSD+TE systems. In addition, teleosts exhibit a wide range of GSD systems, with both classical male (XX/XY) and female heterogamety (ZZ/ZW), but also more complex GSD systems relying on polygenic SD with or without multiple sex chromosomes (Devlin & Nagahama, 2002; Mank & Avise, 2009; Moore & Roberts, 2013). These transitions or turnovers of different GSD systems have been found in closely related species belonging to the same genus (Takehana, Hamaguchi, & Sakaizumi, 2008) and even across populations of the same species (Kallman, 1973). A similar high turnover has also been found for master sex determining (MSD) genes at the top of the genetic sex determination cascade (Matsuda et al., 2002; Myosho et al., 2012; Nanda et al., 2002; Q. Pan et al., 2016; Takehana et al., 2014). Many of these fish MSD genes belong to the “usual suspect” category (Herpin & Schartl, 2015) because they derived from key genes regulating the gonadal sex differentiation network. These “usual suspect” MSD genes currently belong to a few gene families, like the *Dmrt* (Chen et al., 2014; Matsuda et al., 2002; Nanda et al., 2002), *Sox* (Takehana et al., 2014), steroid-pathway (Koyama et al., 2019; Purcell et al., 2018) and Transforming Growth Factor beta (TGFβ) families (Q. Pan et al., 2021), which have been independently and recurrently used to generate new MSD genes. The greatest diversity of MSD genes is found within the TGFβ family with the anti-mullerian hormone, *amh* (Hattori et al., 2012; M. Li et al., 2015; Q. Pan et al., 2019), the gonadal soma derived factor, *gsdf* (Myosho et al., 2012; Rondeau et al., 2013), or the growth/differentiation factor 6, *gdf6* (Imarazene et al., 2021; Reichwald et al., 2015) genes, but also TGF-β type II and type I receptors with the anti-Mullerian hormone receptor type 2, *amhr2* (Feron et al., 2020; Kamiya et al., 2012) and the bone morphogenetic protein receptor, type IBb, *bmpr1bb* (Rafati et al., 2020) genes. However, a few exceptions to the “usual suspects” rule have also been identified with, for instance, the conserved salmonid MSD *sdY* gene that evolved from an immunity-related gene (Bertho, Herpin, Schartl, & Guiguen, 2021; Yano et al., 2012, 2013).

Based on a chromosome-scale high quality genome assembly, and previously published whole-organ transcriptomic data (Pasquier et al., 2016) of *Pangasianodon hypophthalmus*, we identified a male-specific duplication of *amhr2* (*amhr2by*) in that species. This potential Y chromosome-specific *amhr2by* encodes an N-terminal truncated protein that lacks the cysteine-rich extracellular part of the receptor, which is key for proper Amh ligand binding. Sex-linkage studies and genome sequencing of 11 additional Pangasiid species show that *amhr2by* is conserved as a male-specific gene in at least four Pangasiid species, stemming from a single ancient duplication / insertion event at the root of the Siluroidei suborder radiation that is dated around 100 million years ago (Kappas, Vittas, Pantzartzi, Drosopoulou, & Scouras, 2016). Together, these results bring multiple lines of evidence supporting the hypothesis that *amhr2by* is potentially an ancient and conserved master sex-determining gene in Pangasiid catfishes and highlight the recurrent usage of the transforming growth factor β pathway in teleost sex determination.

## MATERIAL AND METHODS

### Samples collection

For high-quality genome reference sequencing, a single *P. hypophthalmus* male was sampled from captive broodstock populations originating from Indonesia and maintained in the experimental facilities of ISEM (Institut des Sciences de l’Evolution de Montpellier, France). High molecular weight (HMW) genomic DNA (gDNA) was extracted from a 0.5-ml blood sample stored in a TNES-Urea lysis buffer (TNES-Urea: 4 M urea; 10 mM Tris-HCl, pH 7.5; 125 mM NaCl; 10 mM EDTA; 1% SDS). HMW gDNA was then purified using a slightly-modified phenol-chloroform extraction (Q. Pan et al., 2019). For the chromosome contact map (Hi-C), 1.5 ml of blood was taken from the same animal and slowly cryopreserved with 15 % dimethyl sulfoxide (DMSO) in a Mr. Frosty Freezing Container (Thermo Scientific) at −80°C. For sex-linkage analyses and short-read genome sequencing, fin clips were sampled and stored in 90% ethanol. *P. djambal* fin clips were sampled from captive broodstock populations originating from Indonesia and maintained in the experimental facilities of ISEM. *P. gigas* fin clips were sampled on broodstock populations kept for a restocking program in Thailand. *P. bocourti* and *P. conchophilus* fin clips were sampled at market places in Vietnam. *P. elongatus*, *P. siamensis*, *P. macronema*, *P. larnaudii*, *P. mekongensis*, and *P. krempfi* were wild samples collected in Vietnam. *P. sanitwongsei* fin clip samples were obtained through the aquaculture trade and their precise origin is unknown.

### Chromosome-scale genome sequencing and assembly of *P. hypophthalmus*

#### Oxford Nanopore sequencing

All library preparations and sequencing were performed using Oxford Nanopore Ligation Sequencing Kits SQK-LSK108 and SQK-LSK109 according to the manufacturer’s instructions (Oxford Nanopore Technologies). For the SQK-LSK108 sequencing Kit, 90 μg of DNA was purified then sheared to 20 kb fragments using the megaruptor1 system (Diagenode). For each library, a DNA-damage repair step was performed on 5 μg of DNA. Then an END-repair-dA-tail step was performed for adapter ligation. Libraries were loaded onto nine R9.4.1 flowcells and sequenced on a GridION instrument at a concentration of 0.1 pmol for 48 h. For the SQK-LSK109 sequencing Kit, 10 μg of DNA was purified then sheared to 20 kb fragments using the megaruptor1 system (Diagenode). For this library, a one-step DNA-damage repair + END-repair-dA-tail procedure was performed on 2 μg of DNA. Adapters were then ligated to DNAs in the library. The library was loaded onto one R9.4.1 flowcell and sequenced on a GridION instrument at a concentration of 0.08 pmol for 48h.

#### 10X Genomics sequencing

The Chromium library was prepared according to 10X Genomics’ protocols using the Genome Reagent Kit v2. The library was prepared from 10 ng of high molecular weight (HMW) gDNA. Briefly, in the microfluidic Genome Chip, a library of Genome Gel Beads, was combined with HMW template gDNA in master mix and partitioning oil to create Gel Bead-In-EMulsions (GEMs) in the Chromium apparatus. Each Gel Bead was then functionalized with millions of copies of a 10x™ barcoded primer. Dissolution of the Genome Gel Bead in the GEM released primers containing (i) an Illumina R1 sequence (Read 1 sequencing primer), (ii) a 16 bp 10x Barcode, and (iii) a 6 bp random primer sequence. The R1 sequence and the 10x™ barcode were added to the molecules during the GEM incubation. P5 and P7 primers, R2 sequence, and Sample Index were added during library construction. 10 cycles of PCR were applied to amplify the library. The library was sequenced on an Illumina HiSeq3000 using a paired-end format with read length of 150 bp with the Illumina HiSeq3000 sequencing kits.

#### Hi-C sequencing

Hi-C library generation was carried out according to a protocol adapted from Rao et al. 2014 (Foissac et al., 2019). The blood sample was spun down, and the cell pellet was resuspended and fixed in 1% formaldehyde. Five million cells were processed for the Hi-C library. After overnight digestion with HindIII (NEB), DNA ends were labeled with Biotin-14-DCTP (Invitrogen) using the klenow (NEB) and religated. In total, 1.4 μg of DNA was sheared to an average size of 550 bp (Covaris). Biotinylated DNA fragments were pulled down using M280 Streptavidin Dynabeads (Invitrogen) and ligated to PE adaptors (Illumina). The Hi-C library was amplified using PE primers (Illumina) with 10 PCR amplification cycles. The library was sequenced using a HiSeq3000 (Illumina, California, USA) in 150 bp paired-end format.

### Genome assembly

GridION data were trimmed using Porechop v0.2.1 (https://github.com/rrwick/Porechop) and filtered using NanoFilt v2.2.0 (De Coster, D’Hert, Schultz, Cruts, & Van Broeckhoven, 2018) with the parameters −1 3000 and -q 7. A d*e novo* assembly was constructed with SmartDeNovo (Ruan, 2015/2019), Wtdbg2 v2.1 (Ruan & Li, 2020) and flye v2.3.7 (Kolmogorov, Yuan, Lin, & Pevzner, 2019), each with default parameters. The resulting assembly metrics were compared, and the draft assembly with the best metrics generated by SmartDeNovo was kept and used as reference. This assembly was then further corrected using long reads. After mapping the trimmed and filtered GridION reads with minimap2 v2.11 (H. Li, 2018) with parameter -x map-ont, the assembly was polished using Racon (Vaser, Sović, Nagarajan, & Šikić, 2017) v1.3.1 with default parameters for three rounds.The assembly was then corrected using short reads. After mapping 10X short reads with Long Ranger v2.1.1, Pilon (Walker et al., 2014) v1.22 was run with parameters --fix bases,gaps -changes. Again, three rounds of these short reads polishing were performed. The final polished genome assembly was then scaffolded using Hi-C information. Reads were aligned to the draft genome using Juicer (Durand, Shamim, et al., 2016) with default parameters. A candidate assembly was then generated with 3D de novo assembly (3D-DNA) pipeline (Dudchenko et al., 2017) with the -r 0 parameter. The candidate assembly was manually reviewed using Juicebox (Durand, Robinson, et al., 2016) assembly tools. Gaps in this chromosome scaled assembly were filled using GapCloser (https://github.com/CAFS-bioinformatics/LR_Gapcloser) v1.1 with default parameters. Reads used to fill these gaps were GridION and PromethION reads filtered with NanoFilt and then corrected with Canu (Koren et al., 2017) v1.6 using parameters –correct genomeSize = 753m –nanopore-raw. The assembly was then corrected one last time using short reads polishing pipeline.

#### Genome analysis and protein-coding gene annotation

K-mer-based estimation of the genome size was carried out with GenomeScope (Vurture et al., 2017) v2.0. 10X reads were processed with Jellyfish v1.1.11 (Marçais & Kingsford, 2011) to count 21-mer with a max k-mer coverage of 10,000 and 1,000,000. BUSCO (Simão, Waterhouse, Ioannidis, Kriventseva, & Zdobnov, 2015) v3.0.2 was run with parameters –species zebrafish and –limit 10 on the single-copy orthologous gene library from the actinopterygii lineage. The first annotation step was to identify repetitive content using RepeatMasker v4.0.7 (https://www.repeatmasker.org/), Dust (Morgulis, Gertz, Schäffer, & Agarwala, 2006), and TRF v4.09 (Benson, 1999). A species-specific *de novo* repeat library was built with RepeatModeler v1.0.11 (http://www.repeatmasker.org/RepeatModeler/) and repeated regions were located using RepeatMasker with the *de novo* and *Danio rerio* libraries. Bedtools v2.26.0 (Quinlan & Hall, 2010) was used to merge repeated regions identified with the three tools and to soft mask the genome. The Maker3 genome annotation pipeline v3.01.02-beta (Holt & Yandell, 2011) combined annotations and evidence from three approaches: similarity with fish proteins, assembled transcripts, and *de novo* gene predictions. Protein sequences from 11 fish species (*Astyanax mexicanus*, *Danio rerio*, *Gadus morhua*, *Gasterosteus aculeatus*, *Lepisosteus oculatus*, *Oreochromis niloticus*, *Oryzias latipes*, *Poecilia formosa*, *Takifugu rubripes*, *Tetraodon nigroviridis*, *Xiphophorus maculatus*) found in Ensembl were aligned to the masked genome using Exonerate v2.4 (Slater & Birney, 2005). RNA-Seq reads of *P. hypophthalmus* (NCBI BioProject PRJNA256973) from the PhyloFish project (Pasquier et al., 2016) were used for genome annotation and aligned to the chromosomal assembly using STAR v2.5.1b (Dobin et al., 2013) with outWigType and outWigStrand options to output signal wiggle files. Cufflinks v2.2.1 (Trapnell et al., 2010) was used to assemble the transcripts that were used as RNA-seq evidence. Braker v2.0.4 (Hoff, Lange, Lomsadze, Borodovsky, & Stanke, 2016) provided *de novo* gene models with wiggle files provided by STAR as hint files for GeneMark (Hoff et al., 2016) and Augustus (Stanke et al., 2006) training. The best supported transcript for each gene was chosen using the quality metric called Annotation Edit Distance (AED) (Eilbeck, Moore, Holt, & Yandell, 2009).

#### miRNA gene and mature miRNA annotation

Small RNA Illumina sequencing libraries were prepared using the NEXTflex Small RNA-Seq Kit v3 (PerkinElmer) following the manufacturer’s instructions and starting with the same total RNA extracts as for the Phylofish project (Pasquier et al., 2016). Total RNA was extracted using Trizol reagent (Euromedex, France) according to the manufacturer’s instructions. Libraries were sequenced on an Illumina HiSeq 2500 sequencer and raw reads were pre-processed using CUTADAPT version 3.4 (Martin, 2011). All eight adult organ libraries (brain, gills, heart ventricle, skeletal muscle, intestine, liver, ovary and testis) were simultaneously analyzed using *Prost!* (Thomas Desvignes, Batzel, Sydes, Eames, & Postlethwait, 2019) selecting for read length 17 to 25 nucleotides and with a minimum of five identical reads. Reads were then aligned to the species’ reference genome using bbmapskimmer.sh version 37.85 of the BBMap suite (https://sourceforge.net/projects/bbmap/). Gene and mature miRNA annotations were performed as previously described (Thomas Desvignes et al., 2019) based on established miRNA gene orthologies among ray-finned fish species (Thomas Desvignes, Sydes, Montfort, Bobe, & Postlethwait, 2021) and using previously published miRNA annotations in spotted gar, zebrafish, three-spined stickleback, Japanese medaka, shortfin molly and blackfin icefish as reference (Braasch et al., 2016; Thomas Desvignes et al., 2019, 2021; Kelley et al., 2021; B.-M. Kim et al., 2019). miRNA and isomiR nomenclature follow the rules established for zebrafish (T. Desvignes et al., 2015).

### Short-read sequencing and genome-guided assemblies of other Pangasiids

#### Short-read sequencing

The *P. gigas* and *P. djambal* genomes were sequenced using an Illumina 2×250 bp format. DNA library construction was performed according to the manufacturer’s instruction using the Truseq DNA nano library prep kit (Illumina). Briefly, gDNA was quantified using the HS dsDNA Assay kit on the Qubit (Invitrogen). 200 ng of gDNA were sonicated on a Bioruptor (Diagenode). Sonicated gDNA was end repaired and size selected on magnetic beads aiming for fragments of an average size of 550 pb. Selected fragments were adenylated on their 3’ ends before ligation of Illumina’s indexed adapters. The library was amplified using 8 PCR cycles and verified on a Fragment Analyzer using the HS NGS fragment kit (Agilent). The library was quantified by qPCR using the KAPA Library quantification kit (Roche, ref. KK4824) and sequenced on half a lane of Hiseq2500 in paired end 2×250nt using the clustering and SBS rapid kit following the manufacturer’s instructions. All other species were sequenced using an Illumina 2×150 bp strategy according to Illumina’s protocols using the Illumina TruSeq Nano DNA HT Library Prep Kit. Briefly, DNA was fragmented by sonication, size selection was performed using SPB beads (kit beads) and adaptors were ligated to be sequenced. Library quality was assessed using an Advanced Analytical Fragment Analyzer and libraries were quantified by qPCR using the Kapa Library Quantification Kit. DNA-seq experiments were performed on one Illumina NovaSeq S4 lane using a paired-end read length of 2×150 bp with the Illumina NovaSeq6000 Reagent Kits.

#### Assembly and annotation

The *P. gigas* and *P. djambal* genomes were assembled from 2×250 bp short reads using the DiscovarDeNovo assembler (https://github.com/bayolau/discovardenovo/) with default parameters. For *P. sanitwongsei*, *P. conchophilus*, *P. bocourti*, *P. larnaudii*, *P. mekongensis*, and *P. krempfi*, 2×150 bp reads were assembled using SPADes v.3.11.1 (Bankevich et al., 2012) and then purged using purge_dups (Guan et al., 2020). The *P. elongatus*, *P. macronema* and *P. siamensis* 2×150 bp short reads were assembled with SPADes v.3.14.1 instead of v.3.11.1 because of a higher individual genome heterozygosity (> 1%), followed by a more stringent purge with Redundans v0.14a (Pryszcz & Gabaldón, 2016). All these species were then assembled into pseudo-chromosomes using a reference-guided strategy and the “query assembled as reference” function from DGenies v1.2.0 (Cabanettes & Klopp, 2018), and the GENO_Phyp_1.0 *P. hypophthalmus* assembly used as a reference. Genes from the NCBI annotation of GENO_Phyp_1.0 were then mapped to chromosome-scale assemblies using Liftoff (Shumate & Salzberg, 2021).

### Species and gene phylogenies

Whole-genome species phylogeny analysis was carried out with protein gene annotation from our 12 Pangasidae species combined with protein sequences from *Ictalurus punctatus* (siluriformes) as a Pangasidae outgroup species. Outgroup species protein sequences were retrieved from Ensembl release 103 (Howe et al., 2021). Orthogroups were identified using OrthoFinder (Emms & Kelly, 2019), followed by multiple sequence alignment of concatenated one-to-one orthologs (n = 8151) using MAFFT version 7.475 (Katoh & Standley, 2013). Species tree inference was performed via IO-TREE 2 (Minh et al., 2020), the latter using a standard non-parametric bootstrap (r = 100).

Gene and protein phylogenetic reconstructions were performed on all *amhr2*/Amhr2 homologous sequences from 28 catfish species along with *amhr2* sequences from *Astyanax mexicanus* (characiformes) and *Electrophorus electricus* (gymnotiformes) as siluriformes outgroups (. Full-length CDS were predicted based on their genomic and protein sequence annotation or retrieved from GenBank (see Table S2 and multi-fasta files of these sequences are publicly available at https://doi.org/10.15454/M3HYAX). To verify the tree topology of *amhr2/*Amhr2 homologs, besides complete protein and cDNA sequences, we also constructed phylogenetic trees with only the first and second codons of the coding sequences (Lemey, 2009). All putative CDS and protein sequences were then aligned using MAFFT (version 7.450) (Katoh & Standley, 2013). Residue-wise confidence scores were computed with GUIDANCE 2 (Sela, Ashkenazy, Katoh, & Pupko, 2015), and only well-aligned residues with confidence scores above 0.99 were retained. Phylogenetic relationships among the *amhr2* sequences were inferred with both maximum-likelihood implemented in IQ-TREE (version 1.6.7) (Minh et al., 2020), and Bayesian methods implemented in Phylobayes (version 4.1) (Lartillot, Lepage, & Blanquart, 2009). More precisely, alignment files from either full-length cDNA, third-codon-removed cDNA, or full-length proteins were used for model selection and tree inference with IQ-TREE (version 1.6.7) (Minh et al., 2020) with 1000 bootstraps and the 1000 SH-like approximate likelihood ratio test for robustness. The same alignment files were run in a Bayesian framework with Phylobayes (version 4.1) (Lartillot et al., 2009) using the CAT-GTR model with default parameters, and two chains were run in parallel for approximately 2000 cycles with the first 500 cycles discarded as burnt-in until the average standard deviation of split frequencies remained ≤ 0.001. The resulting phylogenies were visualized with Figtree (version 1.44).

### Selection analysis on *amhr2* sequences

Selection analysis was performed on the *amhr2* phylogeny using Godon (Davydov, Salamin, & Robinson-Rechavi, 2019). Analyses were performed separately for (a) exons conserved in both *amhr2a* and *amhr2by* (“conserved exons”) and (b) the exon region found only in *amhr2a* (“first exons”). Three codon models were used: M8 (Yang, Nielsen, Goldman, & Pedersen, 2000), M8 with codon gamma rate variation (Davydov et al., 2019), and the branch-site model (Zhang, Nielsen, & Yang, 2005) (conserved exons only). For the branch-site model, the branch leading to the *amhr2by* clade was set as the foreground branch.

### Transcriptome analyses

Reads from *P. hypophthalmus* adult organs and embryos (Pasquier et al., 2016) were mapped on the complete *P. hypophthalmus* reference transcriptome using bwa mem version 0.7.17 (H. Li, 2013). Unique mapped reads were then filtered and a raw count matrix was generated with htseq-count (Anders, Pyl, & Huber, 2015) and normalized using DESeq2 (Love, Huber, & Anders, 2014). Genes of interest were extracted from this complete transcriptome dataset and missing values were replaced by a minimal value (0.1) in the normalized raw count matrix. Hierarchical classification was carried out after log transformation and gene median centering using the cluster 3.0 software (de Hoon, Imoto, Nolan, & Miyano, 2004) with an uncentered correlation similarity metric and an average linkage clustering method.

### Read-coverage analyses around the *amhr2a* and *amhr2by* loci in Pangasiids

To assess whether *amhr2by* is a potential Y specific gene in species for which whole genome sequencing was only obtained from one sample, we computed the read coverage throughout the genome and extracted the read coverage information around the *amhr2a* and *amhr2by* loci. In *P, hypophthalmus*, ONT reads were mapped on its own genome assembly using minimap version 2.11 (H. Li, 2018). In other Pangasiids, Illumina pair-end reads were mapped onto the *P. hypophthalmus* genome assembly using bwa version 0.7.17 (H. Li, 2013), indexed using samtools version 1.8 (H. Li et al., 2009) and sorted by PICARD SortSam. Then a pileup file was generated using samtools mpileup (H. Li et al., 2009) with per-base alignment quality disabled and (-B). Subsequently, a sync file containing the nucleotide composition for each position in the reference was created from the pileup file using popoolation mpileup2sync version 1.201 with a min quality of 20 (-min-qual 20) (Kofler, Pandey, & Schlötterer, 2011). Read depth was then calculated in a 10 kb non-overlapping window using PSASS (version 2.0.0, doi:10.5281/zenodo.2615936).

### Primer design

*P. hypophthalmus amhr2a* and *amhr2by* genes were aligned with bioedit version 7.0.5.3 and specific primers were designed based on this alignment to select highly divergent positions for each paralog. Selected primer sequences were forward: 5’-GGAGTCTATAAACCCGTGGTAGC −3’, and reverse: 5’ - CTATGTCACGCTGAACCTCCAGTGT −3’ (expected amplicon size: 153 bp) for the *amhr2by* gene and forward: 5’-GGAGTCTATAAGCCAGCGGTGGCT −3’, and reverse: 5’-CTATGCCAGAATAACCCTGCAATGC −3’ (expected amplicon size: 142 bp) for the *amhr2a* gene.

### DNA extraction for PCR sex genotyping

DNA from fin clips was extracted using a Chelex-based extraction method. Briefly, a piece of fin clip from each sample was placed into a PCR tube, and then 150 μl 5% Chelex and 20 μl 1 mg/ml proteinase K were added to each tube. Tubes were then vortexed and quickly spun down. After that, samples were incubated for 2 h at 56°C followed by boiling 10 min at 99°C. DNA was then centrifuged at 7500 g for 5 min and diluted to 1:2 with double distilled water. Genotyping PCR reactions were run in 12.5 μl with 1.25 μl JumpStart PCR buffer 10X, 0.125 μl 25 mM dNTP, 0.25 μl 10 μM forward and reverse primers, 8.5 μl ddH_2_O and 2 μl DNA. PCR cycling conditions were: 95°C for 3 min as initial denaturation, then 35 cycles for amplification with denaturation at 95°C for 30 s, annealing at 52°C for 30 s and extension at 72°C for 30 s, and finally another more extension at 72°C for 30 s and hold at 4°C.

## RESULTS

### A high-quality chromosome-scale genome assembly of *P. hypophthalmus*

A high-quality reference genome of a male *P. hypophthalmus* was sequenced using a combination of 10X Linked-Reads, Oxford Nanopore long reads and a chromosome contact map (Hi-C). Its genome size based on the kmer linked-reads distribution was estimated around 810 Mb including, respectively 65% and 35 % of unique and repeated sequences. The heterozygosity level of this *P. hypophthalmus* genome was estimated at around 1.2 %. The integration of all sequencing data provided a genome assembly size of 760 Mb (93% of the kmer estimated size), containing 612 contigs, a scaffold N50 of 26.4 Mb (Table 1) and 99.2% of all sequences anchored onto 30 chromosomes after Hi-C integration (see assembly metrics and comparison with other genome assemblies in Table 1). Combining *de novo* gene predictions, homology to teleost proteins, and evidence from transcripts, 25,076 protein-coding genes were annotated in our male *P. hypophthalmus* reference genome using our in-house genome annotation protocol. Because our *P. hypophthalmus* genome assembly has been derived by NCBI to produce a Reference Sequence (RefSeq) record (GCF_009078355.1) and was annotated by the NCBI Eukaryotic Genome Annotation Pipeline, the NCBI annotation will be used thereafter as reference in the following text. In addition to protein-coding genes, 323 microRNA genes (miRNAs) and 389 mature miRNAs were annotated using Illumina small-RNA sequencing data from a panel of eight organs. Gene and mature miRNA annotations as well as analyzed expression patterns are publicly available on FishmiRNA (http://www.fishmirna.org/) (Thomas Desvignes et al., 2022). This genome-wide miRNA annotation represents the first exhaustive miRNA annotation available for a Pangasiid species.

**Table 1:**
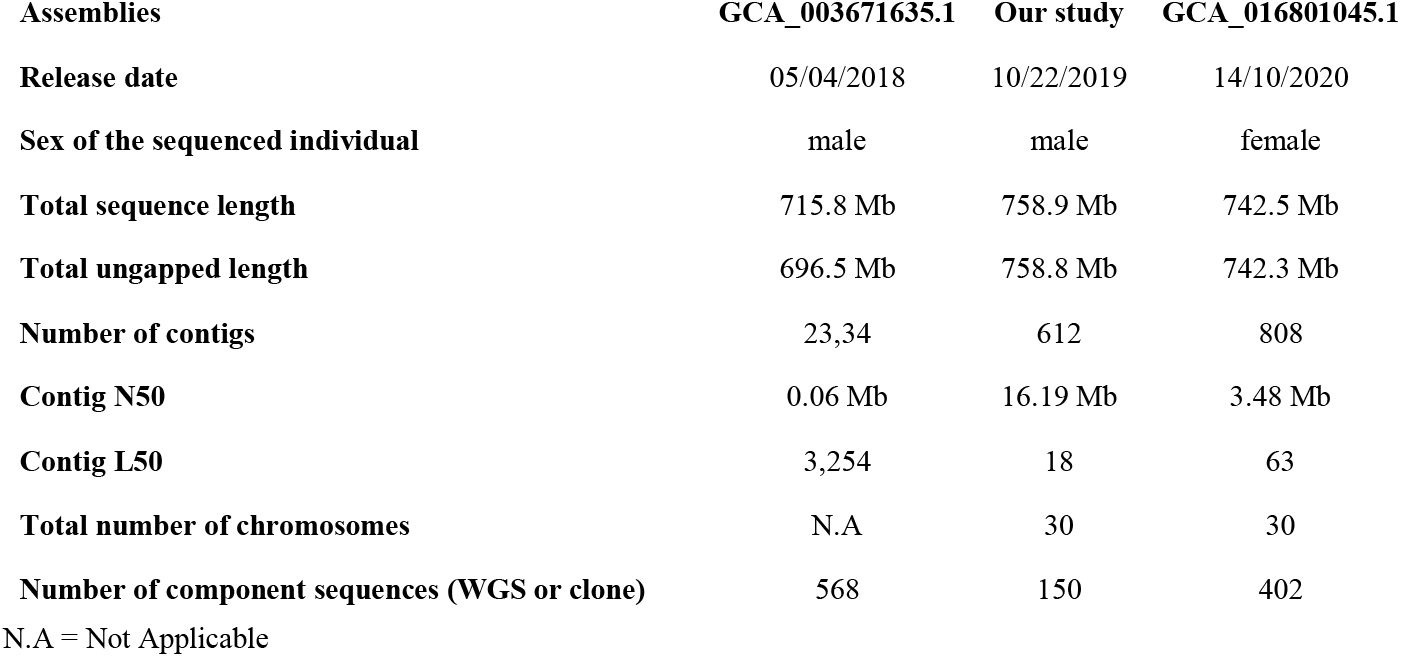
Comparison of our *P. hypophthalmus* reference genome assembly metrics (our study) with the other *P. hypophthalmus* available assemblies.

| Assemblies | GCA_003671635.1 | Our study | GCA_016801045.1 |
| --- | --- | --- | --- |
| Release date | 05/04/2018 | 10/22/2019 | 14/10/2020 |
| Sex of the sequenced individual | male | male | female |
| Total sequence length | 715.8 Mb | 758.9 Mb | 742.5 Mb |
| Total ungapped length | 696.5 Mb | 758.8 Mb | 742.3 Mb |
| Number of contigs | 23,34 | 612 | 808 |
| Contig N50 | 0.06 Mb | 16.19 Mb | 3.48 Mb |
| Contig L50 | 3,254 | 18 | 63 |
| Total number of chromosomes | N.A | 30 | 30 |
| Number of component sequences (WGS or clone) | 568 | 150 | 402 |

N.A = Not Applicable

### Characterization of a male-specific *amhr2* duplication in *P. hypophthalmus*

Because many teleost MSD genes evolved from the duplication of an autosomal “usual suspect” gene, we first searched for potential duplicates of *dmrt1, amh, amhr2, sox3, gsdf* and *gdf6* genes in the *P. hypophthalmus* genome assemblies. We found no gene duplication for *dmrt1, amh, sox3, gsdf* and *gdf6* (*gdf6a* and *gdf6b*), but two *amhr2* homologs were found in the two male *P. hypophthalmus* assemblies (GENO_Phyp_1.0 and VN_pangasius) while only one *amhr2* gene was detected in the female *P. hypophthalmus* ASM1680104v1 assembly. In the GENO_Phyp_1.0 *P. hypophthalmus* assembly, these two *amhr2* homologs, i.e., LOC113540131 (annotated as bone morphogenetic protein receptor type-2-like) and LOC113533735 (annotated as anti-Mullerian hormone type-2 receptor-like) are located respectively on chromosome 4 (Chr04:32,081,919-32,105,291) and 10 (Chr10: 26,334,822-26,348,340). The single *amhr2* locus found in the female ASM1680104v1 assembly (in ASM1680104v1 Chr04), is on chromosome 4 and shares 99% identity over 13.5 kb (100% overlap) with LOC113533735, and 87% identity on only 3% overlapping regions with LOC113540131. Using primers (see Materials and Methods) designed to amplify specifically either LOC113540131 or LOC113533735, we genotyped *P. hypophthalmus* males (N=12) and females (N=11) and found that LOC113540131 is significantly linked with maleness (p = 7.12e^−05^) with a single positive outlier among 11 phenotypic females (see Table 2). In contrast, LOC113533735 was detected in all males and females (Fig. 1). These genotyping results, along with the absence of LOC113540131 in the female ASM1680104v1 assembly, strongly support the hypothesis that LOC113540131 is a Y-specific male-specific, gene. We thus called the LOC113540131 gene, *amhr2by*, as the male-specific Y chromosome paralog of the autosomal LOC113533735 gene named *amhr2a*.

**Table 2:** Sex-linkage of *amhr2by* in five different Pangasiid species. Associations between *amhr2by* specific PCR amplifications and sex phenotypes are provided for both males and females (number of positive individuals for *amhr2by*/total number of individuals) along with the p value of association with sex that was calculated for each species based on the Pearson’s Chi-square test with Yates’ continuity correction.

| Species | males | females | p value |
| --- | --- | --- | --- |
| *Pangasianodon hypophthalmus* | 12/12 | 1/11 | 7.12e-05 |
| *Pangasianodon gigas* | 3/3 | 1/3 | 0.3865* |
| *Pangasius bocourti* | 12/12 | 1/20 | 8.411e-07 |
| *Pangasius conchophilus* | 22/22 | 0/10 | 1.559e-07 |
| *Pangasius djambal* | 6/6 | 0/9 | 8.528e-04 |

\* non-significant association with sex

**Figure 1.**
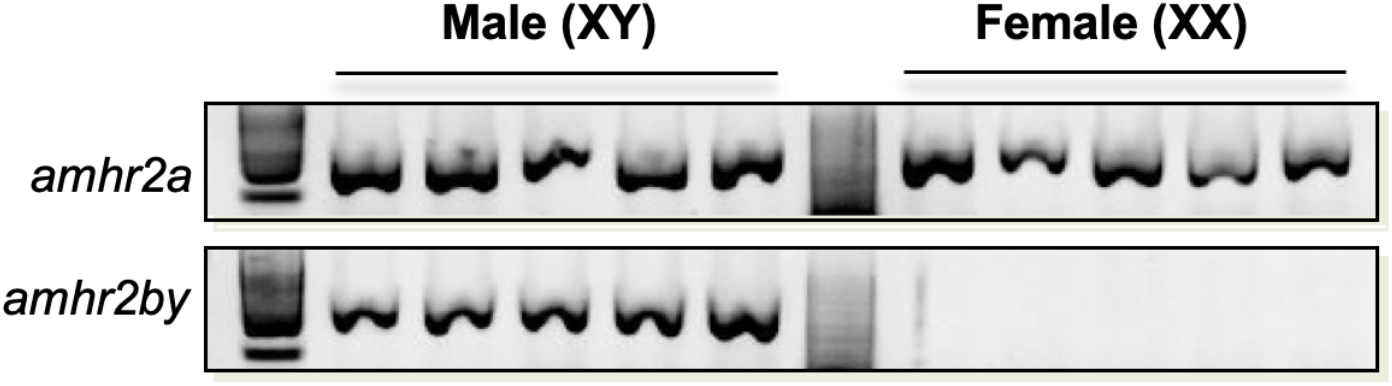
Sex genotyping in *P. hypophthalmus*. The *amhr2a* sequence (upper panel) is PCR amplified in both male and female samples, while the *amh2by* sequence (bottom panel) is only amplified in male samples, indicating that *amhr2by* is male-specific i.e., Y-chromosome linked.

### Comparison of *P. hypophthalmus amhr2by* and *amhr2a* and their inferred proteins

Overall, the predicted structure of the autosomal *P. hypophthalmus amhr2a* and the canonical vertebrate *Amhr2* are similar with the same number of introns and exons. The mVISTA (Frazer, Pachter, Poliakov, Rubin, & Dubchak, 2004) alignments of *P. hypophthalmus amhr2a* and *amhr2by* genes along with their CDS (Fig. 2A), show that these two genes display some sequence identity only within their shared exons, with no significant homology detected in their intronic, 3’UTR, and 5’UTR sequences (Fig. 2A). In addition, the *amhr2by* gene is lacking the first two exons of *amhr2a*, and the third *amhr2by* exon is also truncated. The *amhr2by* and *amhr2a* CDS share 78.78% identity on 1,164 bp of overlapping sequences (78% of the *amhr2a* CDS that is 1,455 bp long). Correspondingly, the two deduced proteins share 70.32% identity over 380 overlapping amino-acids, and Amhr2by lacks 112 amino-acids at its N-terminal extremity corresponding to two first exons and part of exon 3 of Amhr2a. (Fig. 2B and 2C). Hence, the *P. hypophthalmus* Amhr2by translates as an N-terminal-truncated type II receptor lacking its whole extra-cellular domain mediating ligand binding, while overall the remaining of the other functional domains (transmembrane and serine-threonine kinase domain) remain similar between Amhr2a and Amhr2by (Fig. 2B and 2C).

**Figure 2.**
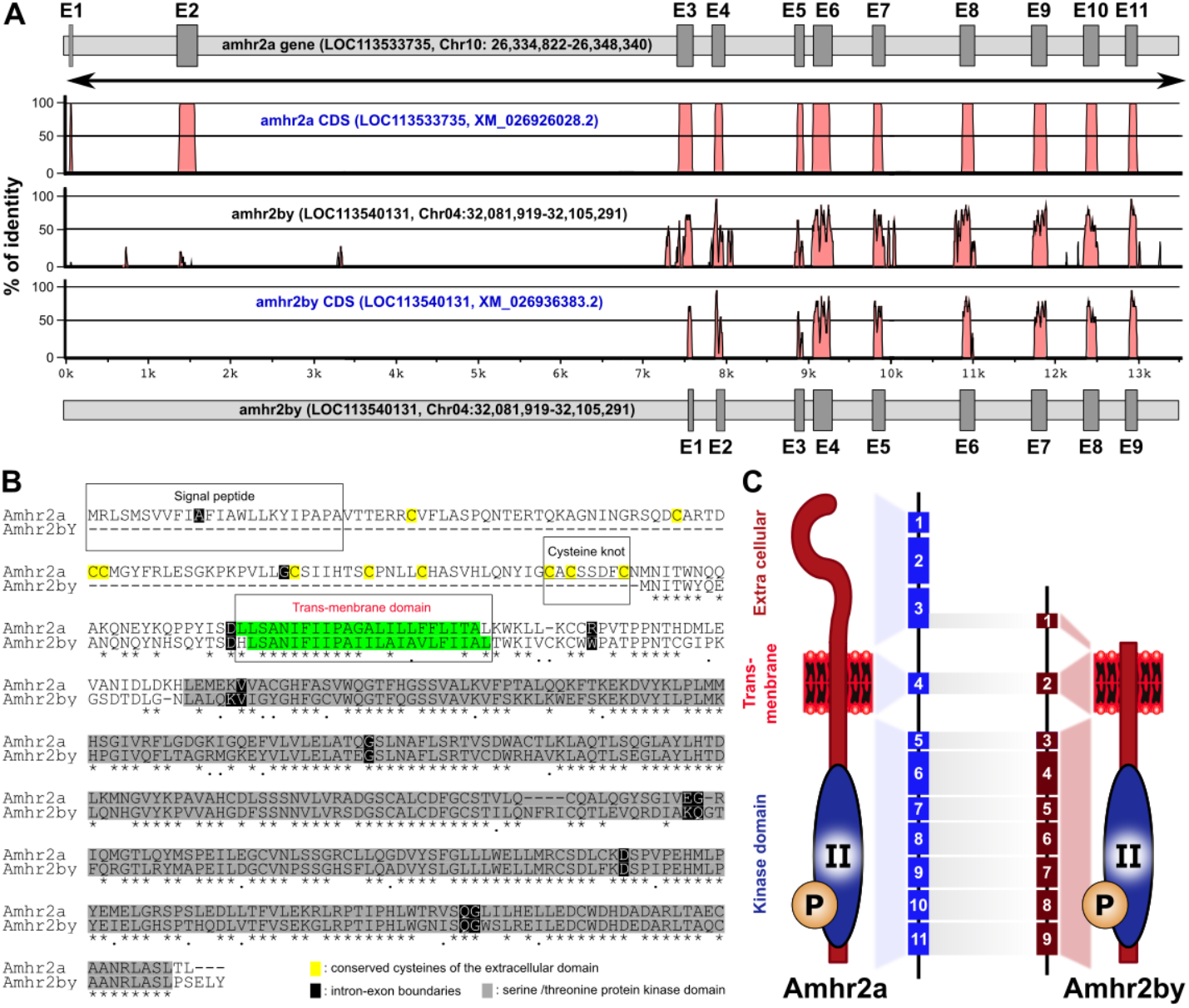
Structure of *amhr2a* and *amhr2by* and deduced proteins in *P. hypophthalmus*. **(A)** Identity plot of the alignment of the autosomal *amhr2a* with the Y-linked *amhr2by* sequences. Exons (E) of both *amhr2* genes are depicted with gray boxes. **(B)** Clustal W alignment of Amhr2a and Amhr2by proteins. Identical amino acids are shaded in gray and conserved cysteines in the extracellular domain of Amhr2a are highlighted in yellow. The different domains (signal peptide, cysteine knot and transmembrane domain) of the receptors are boxed. Intron-exon boundaries are boxed in black for both receptors. **(C)** Schematic representation of *P. hypophthalmus* autosomal Amhr2a and Y-linked Amhr2bY proteins showing the architecture of Amh receptors and the correspondence between exons of Amhr2a and Amhr2by, highlighting the absence of the entire extracellular domain in the truncated Amhr2bY.

### Expression of *amhr2by* and *amhr2a* in *P. hypophthalmus* adult tissues

Using *P. hypophthalmus* RNA-Seq from the PhyloFish database (Pasquier et al., 2016), we examined the organ expression of *amhr2a* and *amhr2by* along with a series of SD genes previously identified in other teleosts, i.e., *amh, dmrt1, gsdf, gdf6a, gdf6b* and *sox3*. Among these genes, *amh, dmrt1*, and *gsdf* display predominant expression in the adult testis and / or ovary with a much lower expression in the eight additional somatic organs examined or in embryos (Fig. 3A). The two *amhr2* genes also have a gonadal-predominant expression pattern with *amhr2a* being expressed in both ovary and testis while *amhr2by* being strictly expressed in the testis as expected for a Y chromosome sex determination gene (Fig. 3B). The two *gdf6* paralogs (*gdf6a, gdf6b*) and *sox3* have no expression or a low expression in gonads and are more expressed in embryos for *sox3* and *gdf6a* or in bones and brain for *sox3*.

**Figure 3.**
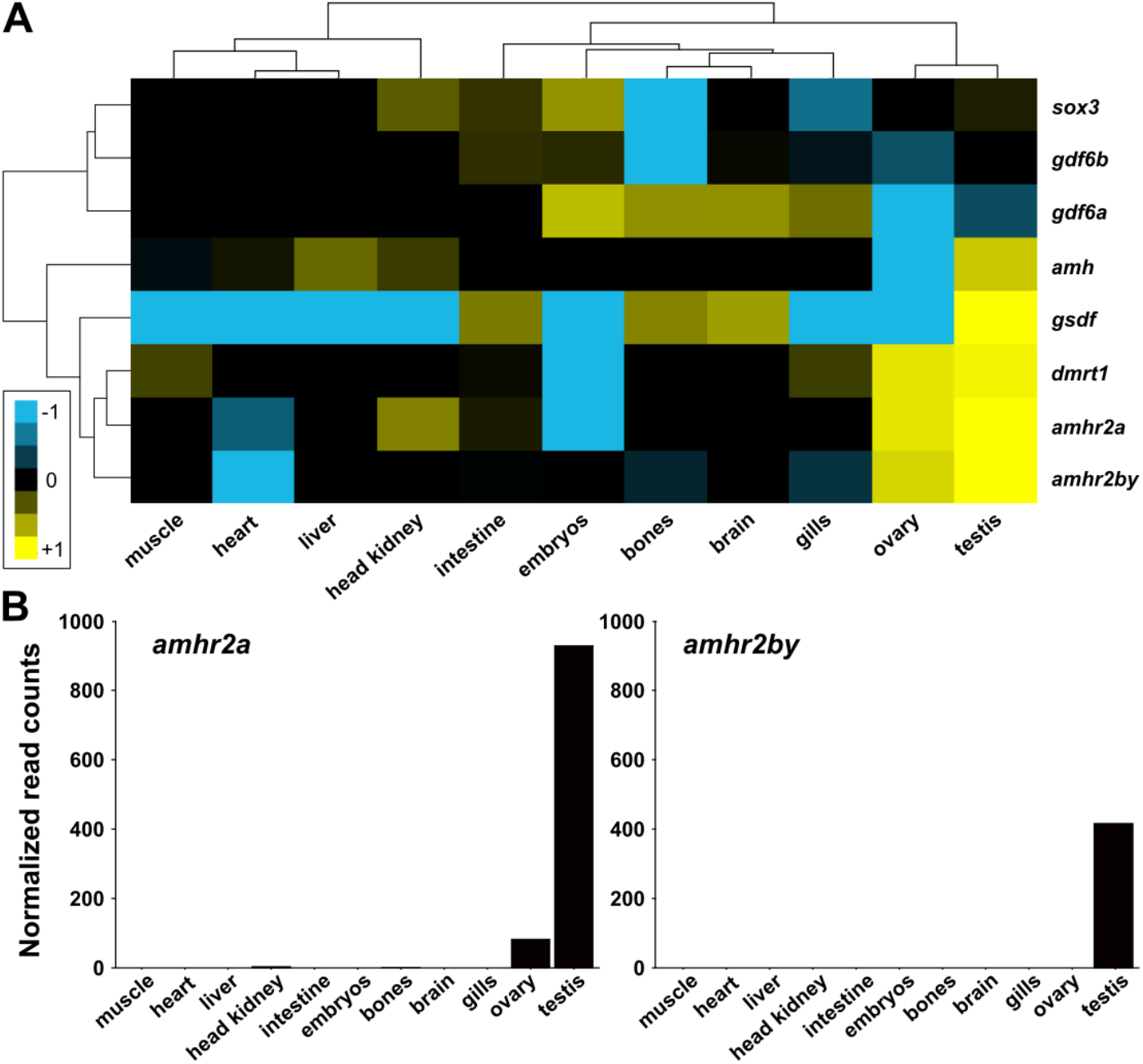
Expression of some sex determination candidate genes in adult organs of *P. hypophthalmus*. **(A)** Hierarchical clustering heatmap analysis of some sex determination genes previously identified in other teleosts, i.e., *amh, amhr2, dmrt1, gsdf, gdf6a, gdf6b* and *sox3* in different organs and embryos of *P. hypophthalmus*. Each colored cell corresponds to a relative expression value (see color legend on the left). **(B)** Normalized read counts of *amhr2a* and *amhr2by* in whole organs and embryos *P. hypophthalmus* transcriptomes.

### Sex-linkage of *amhr2by* in Pangasiids

To explore the evolution of *amhr2by* in Pangasiids, we obtained gDNA samples from 11 additional Pangasiid species with at least some specimens being phenotypically sexed for four of these species (Table S2). Samples from fish that were phenotypically sexed (i.e., *Pangasianodon gigas, Pangasius djambal*, *Pangasius conchophilus*, and *Pangasius bocourti*) were PCR genotyped to explore the potential conservation of *amhr2by* male sex-linkage in Pangasiids. In three of these species, *amhr2by* was found to be significantly associated with male phenotype (p < 8.528e^−04^) (Table 2), except in *P. gigas*, the association was not significant (p = 0.3865) due to the combination of low sample size (3 males and 3 females) and the presence of one female outlier (Table 2). To complement this genotyping information, one male individual of *P. gigas*, *P. djambal*, *P. conchophilus*, and *P. bocourti* and one individual of unknown sex for *P. elongatus*, *P. siamensis*, *P. sanitwongsei*, *P. macronema*, *P. larnaudii*, *P. mekongensis* and *P. krempfi* were sequenced using Illumina short-read strategies. These genomic short-read sequences were assembled and anchored using a reference-guided strategy (Lischer & Shimizu, 2017) on the *P. hypophthalmus* chromosome assembly, and the NCBI gene annotation of GENO_Phyp_1.0 was lifted over to these assemblies (see genome and annotation metrics in Table S1). The *amhr2a* genes were extracted from all these guided assemblies, and *amhr2by* homologs were extracted from the four male assemblies, i.e., *P. gigas, P. djambal, P. conchophilus*, and *P. bocourti* as well as from the unknown sex assemblies of *P. sanitwongsei*, and *P. krempfi*. To better explore sex-linkage in species for which we only sequenced a single individual, read coverage was explored around the *amhr2a* and *amhr2by* loci using the *P. hypophthalmus* as reference genome (Fig. S1). Under the hypothesis that *amhr2by* is also a male-specific Y chromosomal gene in additional Pangasiids, we expected a half coverage around *amhr2by* in males (hemizygous in XY) and an average read coverage around the autosomal *amhr2a*. In agreement with that hypothesis, a half coverage was found around the *amhr2by* locus for all species in which *amhr2by* was identified i.e., the male individuals of *P. hypophthalmus, P.gigas, P. djambal, P. conchophilus*,and *P. bocourti* and individuals of unknown sex in *P. sanitwongsei*, and *P. krempfi*. This result supports hemizygosity of *amhr2by* in these species as expected for a Y chromosomal gene. In other species, i.e., *P. elongatus*, *P. siamensis*, *P. macronema*, *P. larnaudii*, and *P. mekongensis*, no conclusion can be drawn because the absence of finding *amhr2by* in these individuals could be because they are XX females without a Y chromosome and an *amhr2by* gene, or these species may have lost *amhr2by* as a Y chromosome gene.

### Evolution of *amhr2* in Siluriformes

These whole-genome annotations were combined with protein sequences from channel catfish, *Ictalurus punctatus* (Siluriformes, Ictaluridae) used as a Pangasiid outgroup, and 8151 groups of one-to-one orthologs were used after concatenation to construct a whole-genome species tree inference (Fig. 4). In addition, all Pangasiids *amhr2* sequences deduced from our genomic resources were used for phylogenetic analyses with other available catfish *amhr2* genes (Table S2), along with *amhr2* from a gymnotiform (*Electrophorus electricus*) and a characiform (*Astyanax mexicanus*) as the closest species outgroups to the Siluriformes order. The topologies of all trees, i.e., using maximum-likelihood and Bayesian methods on proteins, CDS, and CDS with third codons removed (see Materials and Methods), were all congruent in showing that most of the *amhr2* from the sub-order Siluroidei (Sullivan et al., 2006) cluster with the Pangasiid *amhr2a*, and that outside the Pangasiid family, only a single species (*Pimelodus maculatus*, Pimelodidae) has an *amhr2* duplication clustering with the *amhr2by* sequences (Fig. 5, Fig. S2). Within the Siluriformes, a single *amhr2* in *Corydoras sp* (Callichthyidae, Loricarioidei) roots the *amhr2a* and *amhr2b* duplications (Fig. 5, Fig. S2), suggesting that *amhr2b (P. maculatus*) and *amhr2by* (Pangasiids) arose from an ancient duplication / insertion event at the root of the Siluroidei radiation that is dated around 100 million years ago (Kappas et al., 2016). We also searched for selection acting on the Pangasiid *amhr2* sequences, but detected no statistically significant signal of positive selection (Table 3) for either all exons conserved in both *amhr2a* and *amhr2by* (“conserved exons”) or for the exon region found only in *amhr2a* (“first exons”).

**Figure 4:**
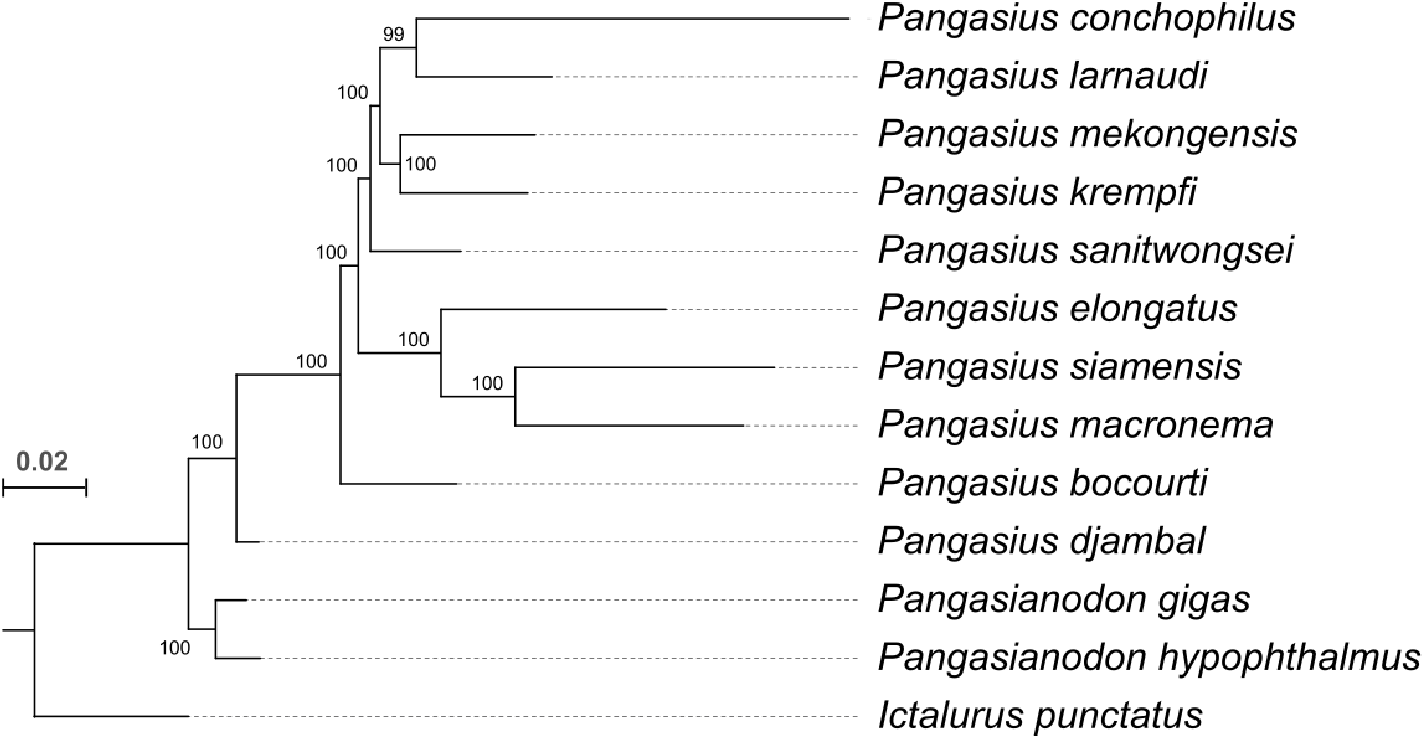
Whole-genome-based phylogenetic tree of all sequenced Pangasiid species. Maximum-likelihood phylogeny of 12 Pangasiidae species with *Ictalurus punctatus* (siluriformes) as a Pangasiidae outgroup, based on alignment of concatenated protein sequences. Branch length scale corresponds to 0.02 amino acid substitutions per site. Support values at each node are proportions of 100 standard non-parametric bootstrap replicates.

**Figure 5.**
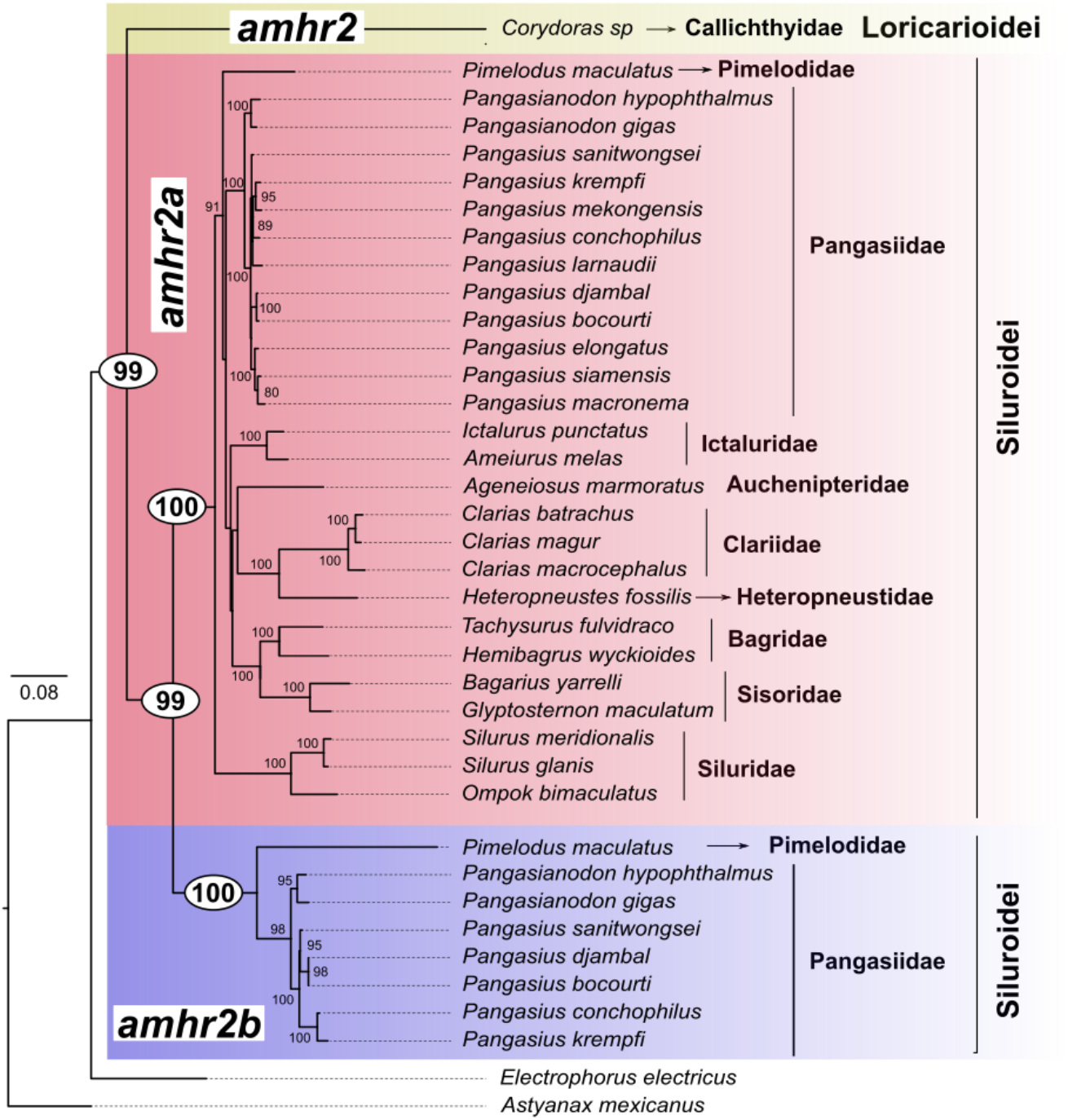
Phylogeny of *amhr2* in catfishes reveals an ancient *amhr2a* / *amhr2b* duplication in Siluroidei. Maximum-likelihood phylogeny of *amhr2* coding sequences (see Supplementary Figure 1 for other phylogenetic approaches) from 28 catfish species with *amhr2* coding sequences from *Astyanax mexicanus* (Characiformes) and *Electrophorus electricus* (Gymnotiformes) as Siluriformes outgroups. Family and suborders are given for all catfish species on the right panel of the figure. The *amhr2b* cluster including the *amhr2by* of Pangasiids is shaded in purple, the *amhr2a* cluster shaded in red, and the *Corydoras sp amhr2* pre-duplication is shaded in yellow. The branch length scale representing the number of substitutions per site is given at the root of the Siluriformes tree. Bootstrap values are given only for values over 80 and are inserted in a white circle at key nodes for the Siluroidei *amhr2* duplication.

**Table 3:** Positive selection analyses reveal no significant signal of positive selection on Pangasiid *amhr2*. P-values were computed using a chi-square distribution with 1 degree of freedom. None of the p-values passed a Bonferroni corrected limit of significance: 0.05/3 = 0.0167. DlnL = difference in log-likelihood between models with and without positive selection; likelihood ratio test statistic.

| Model | Conserved exons |  | First exons |  |
| --- | --- | --- | --- | --- |
|  | DlnL | p-value | DlnL | p-value |
| M8 gamma | 0.0000000 | 0.5000000 | 3.37482 | 3.309990e-02 |
| Branch-site gamma | 0.2976536 | 0.2926786 | - | - |

## DISCUSSION

The Pangasiid family contains both important aquaculture species (Lazard et al., 2009) and key ecological catfish species (Eva et al., 2016) in many south Asian countries. Here, we present a reference genome for striped catfish, *Pangasianodon hypophthalmus*, and provide an additional high-quality genomic resource combining long-read sequencing and a chromosomal assembly for this species. This *de novo* genome (GENO_Phyp_1.0, GCA_009078355.1) was assembled into 30 large scaffolds that most likely correspond to the 30 chromosomes reported previously in cytological studies (Sreeputhorn et al., 2017). This assembly also improves the metrics of the previously publicly available male assembly VN_pangasius (GCA_003671635.1) that was not anchored on chromosomes (O. T. P. Kim et al., 2018), and is comparable in terms of assembly metrics to the newest female ASM1680104v1 (GCA_016801045.1) chromosome-anchored assembly (Z. Gao et al., 2021). In addition to this *P. hypophthalmus* chromosome-anchored assembly, we also provided short-read genome sequencing for eleven additional Pangasiid species belonging to the genera *Pangasianodon* (1 additional species) and *Pangasius* (10 additional species). These short-read assemblies have been anchored and annotated on our reference *P. hypophthalmus* genome assembly and now present a large public set of genomic resources for the Pangasiid family.

Phylogenetic relationships within Siluriformes are still debated with no consensus for clear placement of some families within this order (Kappas et al., 2016; Sullivan et al., 2006). But at a broader scale, it is generally accepted that the sub-order Loricarioidei (defined also as a super-family) containing the armored catfish families (Callichthyids and Loricariids) is the earliest-diverging Siluriformes clade with the Diplomystoidei sub-order being the sister group to the remaining Siluroidei sub-order (Kappas et al., 2016; Sullivan et al., 2006). Pangasiids belong to the Siluroidei sub-order and have been characterized as the sister group to either Ictaluridae and Cranoglanididae (Kappas et al., 2016) or Schilbidae (Villela et al., 2017). Their phylogeny has been explored using both mitochondrial and nuclear makers (Karinthanyakit & Jondeung, 2012; Pouyaud et al., 2016). Here, using a phylogenomic approach (Delsuc, Brinkmann, & Philippe, 2005), we were able to determine the precise phylogenetic relationships among the 12 Pangasiid species for which we produced genome sequencing. Our results confirmed the basal position of the *Pangasianodon* genus as already described (Karinthanyakit & Jondeung, 2012; Na-Nakorn et al., 2006; Pouyaud et al., 2016) and, although we did not sequence any *Helicophagus* or *Pseudolais* genera, results allowed us to resolve the taxonomic positions of several *Pangasius* species (Karinthanyakit & Jondeung, 2012).

The molecular basis of genetic sex determination has been explored in only a few catfishes, with reports on the identification of male sex-specific sequences supporting a XX/XY sex determination system in *Pseudobagrus ussuriensis* (Z.-J. Pan, Li, Zhou, Qiang, & Gui, 2015) and *Pelteobagrus* (*Tachysurus*) *fulvidraco* (Dan, Mei, Wang, & Gui, 2013; Wang, Mao, Chen, Liu, & Gui, 2009) from the Bagridae family, and in *Clarias gariepinus* from the Clariidae family (Kovács, Egedi, Bártfai, & Orbán, 2000). In the Ictalurid channel catfish, *Ictalurus punctatus*, based on whole genome sequencing of a YY individual and genome-wide analyses, an isoform of the breast cancer anti-resistance 1 (*bcar1*) gene has been characterized as the male master sex determining gene (Bao et al., 2019). In Pangasiids, genetic sex-markers have been searched without success in *P. hypophthalmus* and *P. gigas* (Sriphairoj et al., 2007). In our study, based on chromosome-scale genome assemblies of many Pangasiid species, transcriptomic data (Pasquier et al., 2016), and sex-linkage analyses we identified a male-specific duplication of the *amhr2* (*amhr2by*) gene as a potentially conserved male master sex determining gene in that fish family. The role of *Amhr2* as a master sex determining gene has been functionally characterized in the tiger pufferfish, *Takifugu rubripes* and Ayu, *Plecoglossus altivelis* (Kamiya et al., 2012; Nakamoto et al., 2021) and strongly suggested by sex-linkage information in common seadragon, *Phyllopteryx taeniolatus*, alligator pipefish, *Syngnathoides biaculeatus* (Qu et al., 2021), other species of pufferfishes (Duan et al., 2021; F.-X. Gao et al., 2020; Kamiya et al., 2012) and yellow perch, *Perca flavescens* (Feron et al., 2020). In addition, the anti-Mullerian hormone, Amh, which is the cognate ligand of AmhR2, has also been demonstrated or suggested as a master sex determining gene in a few fish species (Hattori et al., 2012; M. Li et al., 2015; Q. Pan et al., 2019, 2021; Song et al., 2021). Our results thus provide a new example of the repeated and independent recruitment of Amh and TGFβ pathway members in fish genetic sex determination (Q. Pan et al., 2021). Although formal proof that this *amhr2by* gene is a conserved master sex determining gene in Pangasiids will require additional gene expression analyses and functional demonstrations, our results `have` application as a useful marker for sex control in many Pangasiid species in aquaculture. Sex dimorphic growth is often one of the main reasons for breeding all-male or all-female populations for aquaculture purposes. In Pangasiids, females have a faster growth rate in *P. djambal* above 3 kg, probably linked with the early maturation of males (Legendre et al., 2000). In contrast, weight gain was better in males compared to females in *P. bocourti* (Meng-Umphan, 2009). In addition, our results will also allow better management of breeders used for restocking in the large and endangered Mekong Giant Catfish, *P. gigas*, because maturation takes as long as 16-20 years in this species (Sriphairoj et al., 2007).

Our results on Pangasiid sex determination also raise interesting questions on Amhr2 structure and evolution. For instance, the N-terminal truncation of all the Pangasiid Amhr2by proteins is intriguing because this N-terminal part of the TGFβ type II receptors encodes the complete extracellular ligand-binding domain that is known to be crucial for ligand binding specificity (Hart et al., 2021). N-terminal truncations of TGFβ receptors acting as sex-determining genes have been already reported for Amhr2 in yellow perch (Feron et al., 2020) and common seadragon (Qu et al., 2021), and for Bmpr1b in the Atlantic herring, *Clupea harengus* (Rafati et al., 2020). In the Atlantic herring, the N-terminal truncated Bmpr1bby protein lacks the canonical TGFβ receptor extracellular domains, but has maintained its ability to propagate a specific intracellular signal through kinase activity and Smad protein phosphorylation (Rafati et al., 2020). Together, these studies suggest that some TGFβ receptors truncated in their N-terminal extracellular ligand-binding domain can still trigger a biological response independent from any ligand activation. The fact that convergently, many fish master sex determining genes encoding a TGFβ receptor with a similar N-terminal truncation, suggests that such a ligand-independent action is probably an important step that could have been selected independently to allow an autonomous action of the master sex determining gene. A second interesting and unexpected result from our study is that the duplication of *amhr2* genes that gave birth to the Pangasiids *amhr2by* gene is potentially ancient and so is likely to still be present in additional catfish species outside the Pangasidae family. This result is well supported by the topologies of our *amhr2* phylogenetic gene trees that place the origin of this duplication at the root of the Siluroidei sub-order that is dated around 100 Mya (Kappas et al., 2016). We also found one example of an *amhr2b* that is retained in the *Pimelodus maculatus* (Pimelodidae family) genome, although we do not know if this gene is also sex-linked in this species. But surprisingly no other *amhr2* duplication has been reported yet in other catfish species. Gains and losses of master sex determining genes have been already described such as in Esociformes in which some species have completely lost the *amh* duplication (*amhby*) that is a master sex determining gene in other closely related species from the same family (Q. Pan et al., 2019, 2021). Such complete gene losses can also be expected in catfishes like for instance in the channel catfish that relies on the *bcar1* gene as master sex determining gene (Bao et al., 2019), with no remains of an *amhr2* gene duplication. This situation is also probably the case for additional catfish species in which we did not find any *amhr2* duplication in male genome assemblies like in the Ictaluridae, *Ameiurus melas*, the Clariidae, *Clarias magur*, and the Auchenipteridae, *Ageneiosus marmoratus*. But if *amhr2b* is also male-specific as in Pangasiids, the question remains open for the additional catfish species where only female genome assemblies are currently available, such as in the Sisoridae, Siluridae and Bagridae families. A more extensive search for a potential duplication of *amhr2* genes in additional Siluroidei catfishes would be needed to better understand the fate of the *amhr2b* gene and whether it remains a master sex determining gene like in the Pangasiid family.

Together our results bring multiple lines of evidence supporting the hypothesis that the conserved Pangasiid *amhr2by* is a potential sex determining gene that stemmed from an ancient duplication common to all Siluroidei catfishes. Our results highlight the recurrent usage of the TGFβ pathway in teleost sex determination (Q. Pan et al., 2021) and the potential functional innovation through protein truncation. Furthermore, our results showcase the less considered long-term stability of sex determination gene in teleosts, a group that often receives attention for its dynamic evolution of sex determination systems.

## DATA AVAILABILITY

The Whole Genome Shotgun project of *P. hypophthalmus*, is available in the Sequence Read Archive (SRA), under BioProject reference PRJNA547555 with 10X genomics and Hi-C Illumina sequencing data is available in SRA under accession number SRX6071341 and SRX6071345 and Oxford Nanopore long reads data under SRA accession numbers SRX6071342 to SRX6071344 and SRX6071346 to SRX6071355. *P. hypophthalmus* small RNA-Seq sequences are available in SRA under Bioproject PRJNA256963. *P. gigas* and *P. djambal* genomes assembled with a *P. hypophthalmus* reference-guided strategy have been submitted to SRA under the respective BioProjects PRJNA593917 and PRJNA605300. All other Pangasiidae genomes assembled with a *P. hypophthalmus* reference-guided strategy without their genome annotations are available in SRA under BioProject PRJNA795327, and their genome assemblies plus their annotations are available in the omics dataverse (Open source research data repository) server with the following DOI (https://doi.org/10.15454/M3HYAX). *Pangasius siamensis* has been considered by NCBI curators as a *P. macronema* synonym and its genome is then recorded in NCBI with *P. macronema* as a Biosample species name, with sample name PaSia (for *Pangasius siamensis*) under accession BioSample number SAMN24707637.

## BENEFIT-SHARING STATEMENT

A research collaboration was developed with scientists from the countries providing genetic samples (KS in Thailand, GR in Indonesia, TTTH in Vietnam, and JR and FLA in Brazil), all collaborators are included as co-authors, the results of research have been shared with the provider communities, and the research addresses a priority concern, in this case the conservation of organisms being studied. More broadly, our group is committed to international scientific partnerships, as well as institutional capacity building.

## ACKNOWLEDGEMENTS

This project was supported by funds from the “Agence Nationale de la Recherche” (PhyloSex project, grant No. ANR-13-ISV7-0005; GenoFish project, grant No. ANR-16-CE12-0035) and R35 GM139635 and R01 OD011116 from the National Institutes of Health, USA. The GeT-PlaGe, Gentyane and Montpellier GenomiX (MGX) facilities were supported by France Génomique National infrastructure, funded as part of “Investissement d’avenir” program managed by Agence Nationale pour la Recherche (grant No. ANR-10-INBS-09). MRR was supported by the Swiss National Science Foundation (grant 173048). We are grateful to the Genotoul bioinformatics platform Toulouse Midi-Pyrenees (Bioinfo Genotoul) for providing computing and/or storage resources.

## AUTHOR CONTRIBUTIONS

YG, and JHP designed the project. JCA, RD, MC, TTTH, RG, KS, JR and FLA collected the samples, EJ, MW, CI, AC, CR, OB, SV, CL, CP, EB, VG and HA extracted the gDNA, made the genomic libraries and sequenced them. CC, CK, MZ, MW, QP and YG processed the genome assemblies and / or analyzed the results. TD, JM and JB processed and analyzed the small RNA sequencing data for miRNA analysis. CFB, MW, QP and MRR performed phylogenetic analyses. CFB and MRR performed the selection analysis. MW, JHP, CC, CK, CR, QP and YG wrote the manuscript with inputs from all other coauthors. JHP, CD, JB and YG, supervised the project administration and raised funding. All the authors read and approved the final manuscript.

---

**Supplementary Figure 1.**
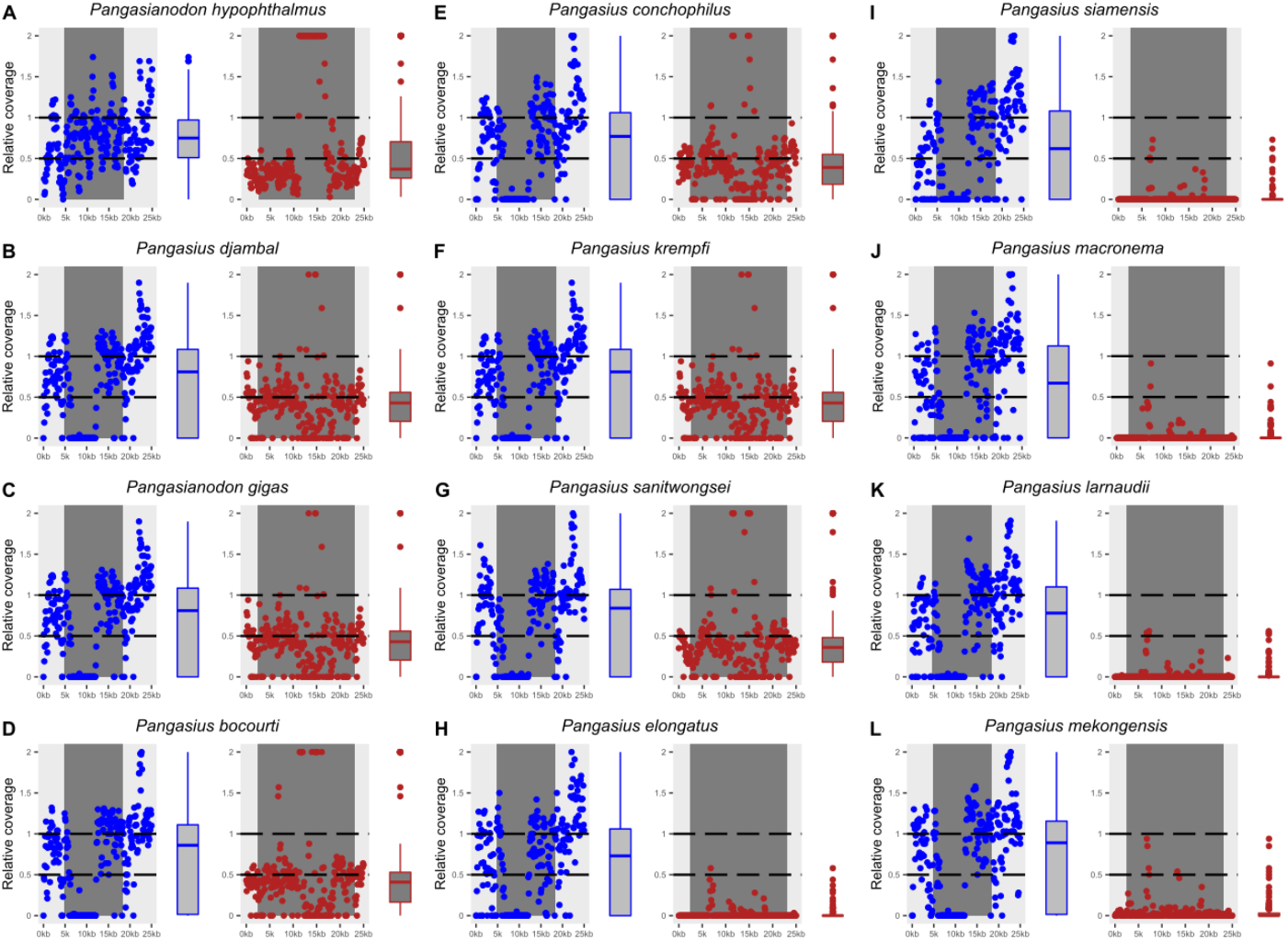
Relative genome read coverage around the *amhr2a* and *amhr2by* loci in 12 Pangasiids supports hemizygosity of *amhr2by* in some species. **(A-L)** Relative average read coverage was deduced from each species short-read remapping on the *P. hypophthalmus* genome reference and is shown in blue for the autosomal *amhr2a* locus and in red for the *amhr2by* locus (left and right side respectively of each species panel). Half read coverage compared to genome average was detected around the *amhr2by* locus compared to the *amhr2a* locus in the male genomes of *P. hypophthalmus, P. gigas*, *P. djambal*, *P. conchophilus*, and *P. bocourti* (**A-E**) and in the unknown sex genomes of *P. sanitwongsei*, and *P. krempfi* (**F-G**), supporting hemizygosity of *amhr2by* in these species as it would be expected for a Y chromosomal gene. In *P. elongatus*, *P. siamensis*, *P. macronema*, *P. larnaudii*, and *P. mekongensis* (**H-L**) no reads were significantly remapped on the *P. hypophthalmus amhr2by* locus either because these sequenced individuals are XX females without a Y chromosome *amhr2by* gene, or because these species have lost *amhr2by* as a Y chromosome gene.

**Supplementary Figure 2.**
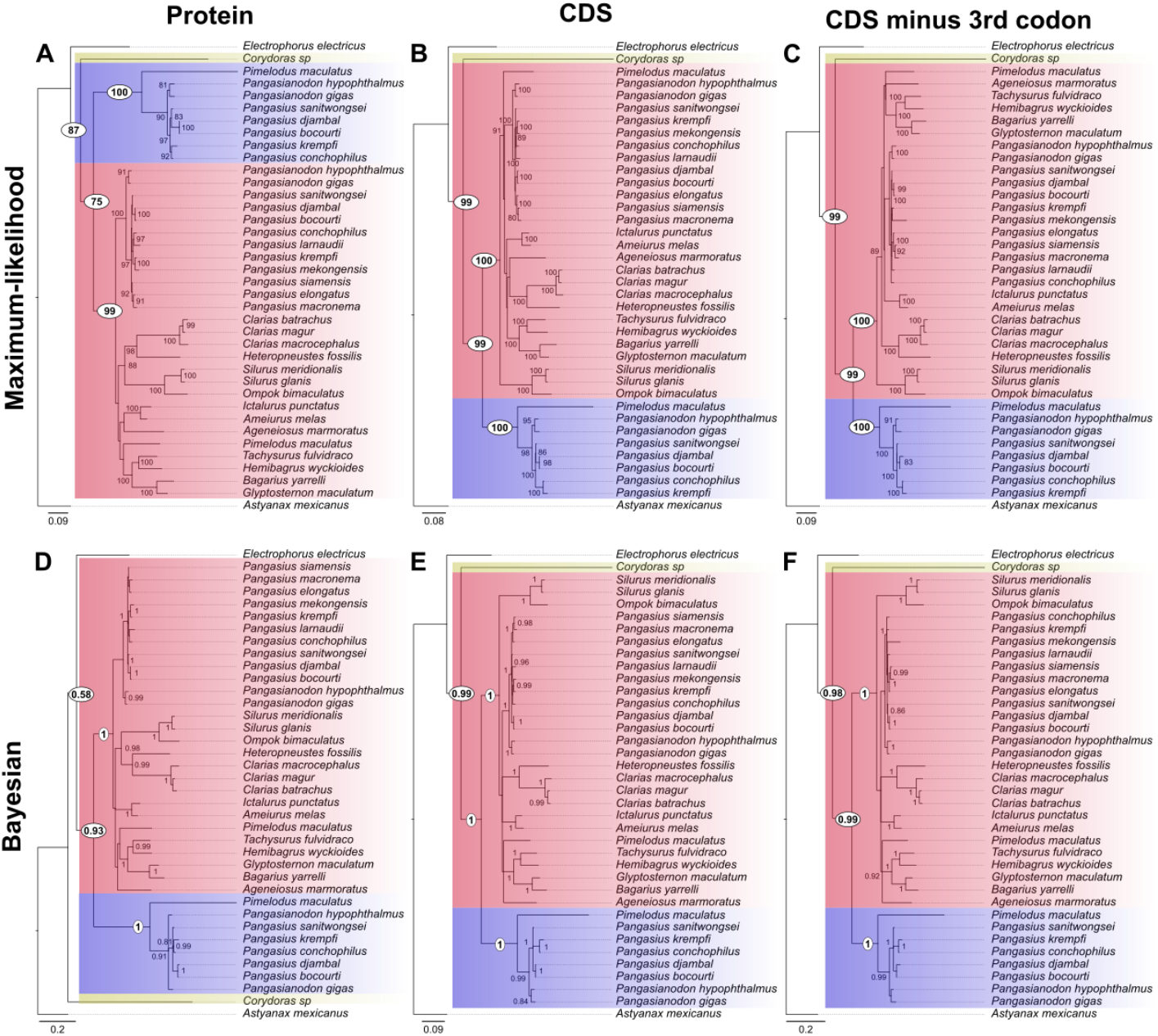
Phylogenies of Amhr2 / *amhr2* in catfishes support an ancient *amhr2a* / *amhr2b* duplication in Siluroidei. Maximum-likelihood (**A**, **B**, **C**) and Bayesian (**D**, **E**, **F**) phylogenies of Amhr2 proteins (**A**, **D**), *amhr2* coding (CDS) sequences (**B**, **E**) and *amhr2* CDS sequences with the third codon removed (**C**, **F**) from 28 catfish species with sequences from *Astyanax mexicanus* (Characiformes) and *Electrophorus electricus* (Gymnotiformes) as Siluriformes outgroups. The *amhr2b* cluster including the *amhr2by* of Pangasiids is shaded in purple, the *amhr2a* cluster shaded in red, and the *Corydoras sp amhr2* pre-duplication is shaded in yellow. The branch length scale representing the number of substitutions per site is given below each tree. Bootstrap values are given only for values over 80 except at key nodes for the Siluroidei *amhr2* duplication (white circles).

**Supplementary Table 1:** Genome assembly characteristics and annotation metrics of 12 Pangasiid species.

| Species | Sex | Sequencing/assembly | Guided assembly | N | G.S (Gb) | Max (Mb) | N50 (Mb) | L50 | % Chr | Annotation | Buscos (C) | Buscos (S) | Buscos (D) | Buscos (F) | Buscos (M) |
| --- | --- | --- | --- | --- | --- | --- | --- | --- | --- | --- | --- | --- | --- | --- | --- |
| *Pangasianodon hypophthalmus* | male | ONT, 10X, Hi-C<br>Smartdenovo/lonranger/juicer | N.A | 612 | 0.759 | 35.6 | 26.16 | 13 | 99.2 | de novo / GenBank | 96.6 | 95.6 | 1.0 | 1.0 | 2.4 |
| *Pangasianodon gigas* | male | Illumina 2x 250 bp<br>Discovar de novo | Dgenies | 283151 | 0.841 | 35.47 | 26.7 | 12 | 89.0 | de novo /<br>in house |  |  |  |  |  |
| *Pangasius djambal* | male | Illumina 2x 250 bp<br>Discovar de novo | Dgenies | 415588 | 0.867 | 34.66 | 28.07 | 11 | 82.7 | de novo /<br>in house |  |  |  |  |  |
| *Pangasius conchophilus* | male | Illumina 2x150 bp<br>SPADes v.3.11.1 /purge_dup | Dgenies | 937794 | 0.815 | 30.1 | 17.57 | 18 | 73.0 | Lifted from<br>*P. hypophthalmus* | 81.9 | 80.7 | 1.2 | 6.2 | 11.9 |
| *Pangasius bocourti* | male | Illumina 2x150 bp<br>SPADes v.3.11.1 /purge_dup | Dgenies | 740730 | 0.780 | 33.3 | 24.19 | 14 | 86.7 | Lifted from<br>*P. hypophthalmus* | 95.5 | 94.6 | 0.9 | 1.4 | 3.1 |
| *Pangasius elongatus* | U | Illumina 2x150 bp<br>SPADes v.3.14.1 /redundans v0.14a | Dgenies | 126560 | 0.712 | 31.0 | 22.43 | 14 | 87.6 | Lifted from<br>*P. hypophthalmus* | 90.7 | 89.8 | 0.9 | 3.1 | 6.2 |
| *Pangasius siamensis* | U | Illumina 2x150 bp<br>SPADes v.3.14.1 /redundans v0.14a | Dgenies | 53159 | 0.685 | 32.4 | 23.59 | 13 | 95.4 | Lifted from<br>*P. hypophthalmus* | 93.0 | 92.0 | 1.0 | 2.1 | 4.9 |
| *Pangasius sanitwongsei* | U | Illumina 2x150 bp<br>SPADes v.3.11.1 /purge_dup | Dgenies | 387468 | 0.743 | 34.1 | 24.55 | 14 | 92.4 | Lifted from<br>*P. hypophthalmus* | 96.3 | 95.6 | 0.7 | 1.1 | 2.6 |
| *Pangasius macronema* | U | Illumina 2x150 bp<br>SPADes v.3.14.1 /redundans v0.14a | Dgenies | 42101 | 0.683 | 32.4 | 23.73 | 13 | 95.9 | Lifted from<br>*P. hypophthalmus* | 93.2 | 91.9 | 1.3 | 2.3 | 4.5 |
| *Pangasius larnaudii* | U | Illumina 2x150 bp<br>SPADes v.3.11.1 /purge_dup | Dgenies | 375123 | 0.733 | 33.0 | 23.84 | 14 | 91.3 | Lifted from<br>*P. hypophthalmus* | 95.6 | 94.8 | 0.8 | 1.5 | 2.9 |
| *Pangasius mekongensis* | U | Illumina 2x150 bp<br>SPADes v.3.11.1 /purge_dup | Dgenies | 443231 | 0.766 | 33.7 | 24.28 | 14 | 88.8 | Lifted from<br>*P. hypophthalmus* | 93.6 | 92.8 | 0.8 | 2.5 | 3.9 |
| *Pangasius krempfi* | U | Illumina 2x150 bp<br>SPADes v.3.11.1 /purge_dup | Dgenies | 448669 | 0.739 | 33.5 | 24.31 | 14 | 91.3 | Lifted from<br>*P. hypophthalmus* | 95.1 | 94.2 | 0.9 | 1.8 | 3.1 |

Sex = phenotypic sex of the animal sequenced (U= unknown), N= Number of contigs, G.S = genome assembly size (kb), Max = size of the longest scaffold, N50 = scaffold N50 (Mb), L50 = scaffold L50, % Chr = percentage of the assembly in chromosomes, Buscos (V4, in genome mode with actinopterygii lineage) score in percentage (C = Complete, S = Single copy, D = Duplicated, F = Fragmented, M = Missing). N.A = Not Applicable.

**Supplementary Table 2:** Origin of the catfish *amhr2* sequences used for phylogenetic analyses.

| Species | Family | Sub-order | order | Sex | Gene | Source | Sequences deduced from |
| --- | --- | --- | --- | --- | --- | --- | --- |
| *Pangasianodon hypophthalmus* | Pangasiidae | Siluroidei | Siluriformes | male | *amhr2a*<br>*amhr2by* | This study | Genome annotation |
| *Pangasianodon gigas* | Pangasiidae | Siluroidei | Siluriformes | male | *amhr2a*<br>*amhr2by* | This study | Genome annotation |
| *Pangasius djambal* | Pangasiidae | Siluroidei | Siluriformes | male | *amhr2a*<br>*amhr2by* | This study | Genome annotation |
| *Pangasius conchophilus* | Pangasiidae | Siluroidei | Siluriformes | male | *amhr2a*<br>*amhr2by* | This study | Genome annotation |
| *Pangasius bocourti* | Pangasiidae | Siluroidei | Siluriformes | male | *amhr2a*<br>*amhr2by* | This study | Genome annotation |
| *Pangasius elongatus* | Pangasiidae | Siluroidei | Siluriformes | unknown | *amhr2a* | This study | Genome annotation |
| *Pangasius siamensis* | Pangasiidae | Siluroidei | Siluriformes | unknown | *amhr2a* | This study | Genome annotation |
| *Pangasius sanitwongsei* | Pangasiidae | Siluroidei | Siluriformes | unknown | *amhr2a*<br>*amhr2by* | This study | Genome annotation |
| *Pangasius macronema* | Pangasiidae | Siluroidei | Siluriformes | unknown | *amhr2a* | This study | Genome annotation |
| *Pangasius larnaudii* | Pangasiidae | Siluroidei | Siluriformes | unknown | *amhr2a* | This study | Genome annotation |
| *Pangasius mekongensis* | Pangasiidae | Siluroidei | Siluriformes | unknown | *amhr2a* | This study | Genome annotation |
| *Pangasius krempfi* | Pangasiidae | Siluroidei | Siluriformes | unknown | *amhr2a*<br>*amhr2by* | This study | Genome annotation |
| *Clarias batrachus* | Clariidae | Siluroidei | Siluriformes | unknown | *amhr2a* | GCA_003987875.1 | inferred from genome assembly |
| *Clarias macrocephalus* | Clariidae | Siluroidei | Siluriformes | female | *amhr2a* | GCA_011419295.1 | inferred from genome assembly |
| *Clarias magur* | Clariidae | Siluroidei | Siluriformes | male | *amhr2a* | GCA_013621035.1 | inferred from genome assembly |
| *Bagarius yarrelli* | Sisoridae | Siluroidei | Siluriformes | female | *amhr2a* | GCA_005784505.1 | inferred from genome assembly |
| *Glyptosternon maculatum* | Sisoridae | Siluroidei | Siluriformes | female | *amhr2a* | http://gigadb.org/dataset/view/id/100489 | inferred from genome assembly |
| *Silurus glanis* | Siluridae | Siluroidei | Siluriformes | female | *amhr2a* | GCA_014706435.1 | inferred from genome assembly |
| *Silurus meridionalis* | Siluridae | Siluroidei | Siluriformes | female | *amhr2a* | GCA_014805685.1 | KAF7704051.1 |
| *Ompok bimaculatus* | Siluridae | Siluroidei | Siluriformes | unknown | *amhr2a* | GCA_009108245.1 | inferred from genome assembly |
| *Hemibagrus wyckioides* | Bagridae | Siluroidei | Siluriformes | female | *amhr2a* | GCA_019097595.1 | KAG7327988.1 |
| *Tachysurus fulvidraco* | Bagridae | Siluroidei | Siluriformes | female | *amhr2a* | GCF_003724035.1 | XP_027015428.1 |
| *Ageneiosus marmoratus* | Auchenipteridae | Siluroidei | Siluriformes | male | *amhr2a* | GCA_003347165.1 | inferred from genome assembly |
| *Heteropneustes fossilis* | Heteropneustidae | Siluroidei | Siluriformes | male | *amhr2a* | unpublished | Inferred from transcriptome information |
| *Ameiurus melas* | Ictaluridae | Siluroidei | Siluriformes | male | *amhr2a* | GCA_012411365.1 | KAF4083677.1 |
| *Ictalurus punctatus* | Ictaluridae | Siluroidei | Siluriformes | female | *amhr2a* | GCF_001660625.1 | XP_017331275.1 |
| *Pimelodus maculatus* | Pimelodidae | Siluroidei | Siluriformes | male | *amhr2a*<br>*amhr2b* | unpublished | inferred from genome assembly |
| *Corydoras sp C115* | Callichthyidae | Loricarioidei | Siluriformes | unknown | *amhr2* | GCA_019802505.1 | inferred from genome assembly |
| *Electrophorus electricus* | Gymnotidae | NR | Gymnotiformes | unknown | *amhr2* | GCF_013358815.1 | XP_035376390.1 |
| *Astyanax mexicanus* | Stethaprioninae | NR | Characiformes | female | *amhr2* | GCF_000372685.2 | XP_022538368.1 |

NR : not relevant

